# OSBA: An open neonatal neuroimaging atlas and template for spina bifida aperta

**DOI:** 10.1101/2024.06.04.597318

**Authors:** Anna Speckert, Hui Ji, Kelly Payette, Patrice Grehten, Raimund Kottke, Samuel Ackermann, Beth Padden, Luca Mazzone, Ueli Moehrlen, SPINA BIFIDA STUDY GROUP ZURICH, Andras Jakab

**Affiliations:** Center for MR-Research, University Children’s Hospital Zurich, Zurich, Switzerland; University Research Priority Program (URPP), Adaptive Brain Circuits in Development and Learning (AdaBD), University of Zurich; Department of Diagnostic Imaging, University Children’s Hospital Zurich; The Zurich Center for Fetal Diagnosis and Therapy, Switzerland; Division of Pediatric Rehabilitation, University Children’s Hospital Zurich; Zurich Center for Spina Bifida, University Children’s Hospital Zurich; Department of Pediatric Surgery, University Children’s Hospital Zurich; Faculty of Medicine, University of Zurich; Spina Bifida Study Group Zurich, Switzerland

**Keywords:** spina bifida aperta, neonatal atlas, T2-weighted MRI, diffusion MRI

## Abstract

We present the Open Spina Bifida Aperta (OSBA) atlas, an open atlas and set of neuroimaging templates for spina bifida aperta (SBA). Traditional brain atlases may not adequately capture anatomical variations present in pediatric or disease-specific cohorts. The OSBA atlas fills in this gap by representing the computationally averaged anatomy of the neonatal brain with SBA after fetal surgical repair. The OSBA atlas was constructed using structural T2-weighted and diffusion tensor MRI of 28 newborns with SBA who underwent prenatal surgical correction. The corrected gestational age at MRI was 38.1 ± 1.1 weeks (mean ± SD). The OSBA atlas consists of T2-weighted and fractional anisotropy templates, along with 9 tissue prior maps and region of interest (ROI) delineations. The OSBA atlas offers a standardized reference space for spatial normalization and anatomical ROI definition. Our image segmentation and cortical ribbon definition is based on a human-in-the-loop approach including manual segmentation. The precise alignment of the ROIs was achieved by a combination of manual image alignment and automated, non-linear image registration. By providing a dedicated neonatal atlas for SBA, we enable more accurate spatial standardization and support advanced analyses such as diffusion tractography and connectomic studies in newborns affected by this condition.

## 1. Summary

Neuroimaging atlases are indispensable for image analysis. They consist of templates which represent common features of the human brain by averaging the brain anatomy of a population [1], as well as maps representing semantic neuroanatomical labels. In the field of neuroimaging analysis, the most common source for constructing atlases are MR images from a group of individuals representative of the group the atlas is meant to represent. These images are then spatially normalized to a common coordinate system through registration and fusion methods. Atlases are particularly useful for carrying out group-level measurements in a subject cohort after spatial normalization, measuring variability in brain anatomy, and establishing localization in functional experiments. Atlases are particularly useful for multicenter studies [1,2].

However, atlases designed for the general population might not be suitable for specific groups, such as paediatric or disease-specific populations [3]. This has led to the development of more tailored atlases for such populations [3]. This remains a challenge for the developing brain, where extensive anatomical changes take place in a short period of time, requiring age-specific atlases [4,5]. Having only age-specific atlases is insufficient when fetal or neonatal brain anatomy diverges from normal development, or for diseases such as open spina bifida, or spina bifida aperta (SBA), one of the most prevalent fetal abnormalities affecting the central nervous system [6]. Fidon [2] confronted this challenge and developed a spatio-temporal fetal brain atlas for SBA.

SBA manifests when the neural tube does not successfully close within the first four weeks following conception [2]. SBA manifests in multiple brain malformations, most frequently abnormal corpus callosum, hypoplastic pons, enlargement of the ventricles, cerebellar malformation, and hypoplastic mesencephalon [7]. Spatially normalizing MR images for neuroimaging analysis is particularly challenging when dealing with subjects whose anatomical structures are underdeveloped, malformed, or enlarged to varying degrees, as accurately establishing correspondences becomes inherently difficult in these cases. Not only is the fetal period of SBA crucial to study, but so is the neonatal phase. The most frequently reported anomalies during this stage, as highlighted by Mufti’s [8] review, are callosal dysgenesis and heterotopia. These anomalies are likely linked to disruptions in neural migration, influenced by the altered CSF dynamics in SBA [9]. The challenge when studying neonatal SBA brains lies in the lack of temporal and disease-specific atlases which could provide anatomical context to the MR images. While there are neonatal atlases available for typically developing brains, such as the Edinburgh Neonatal Atlas (ENA33), which parcellates the brain into 107 anatomical regions and was created through temporal backpropagation of the widely used adult brain atlas (SRI24/TZO) to the neonatal date [10], dedicated neonatal atlases for SBA analysis are currently unavailable. Relying on healthy neonatal atlases for SBA analysis poses a challenge due to ambiguous mapping between regions of interest (ROI) to SBA brains, marked by heterotopia, callosal dysgenesis and enlarged ventricles.

Here we present a new, open atlas and set of neuroimaging templates for SBA, grounded in the ENA33 atlas, enabling direct comparisons, and supporting future research on neonates with SBA. The Open Spina Bifida Aperta (OSBA) atlas addresses this gap, namely, a need for a dedicated neonatal SBA atlas that ideally aligns with those designed for healthy neonatal brains, facilitating case-control studies. The OSBA atlas was constructed using widely-used normalization and atlas creation tools and comprise of T2-weighted and diffusion tensor MRI-based templates of newborns with SBA. The anatomical label maps comprise of tissue prior maps and ROIs according to the definition of the ENA33 atlas. The OSBA atlas is a valuable tool for spatial standardization in SBA and for a wide range of applications where ROIs are necessary for defining anatomical regions for between-subject statistical analysis. An example is studying the connectivity structure of the newborn brain with SBA. In such studies, the OSBA atlas would provide a joint reference space for spatial normalization as well as ROIs to define the seeds for diffusion tractography and consequently, the nodes for connectomic analysis.

## 2. Data Description

**Table 1.**
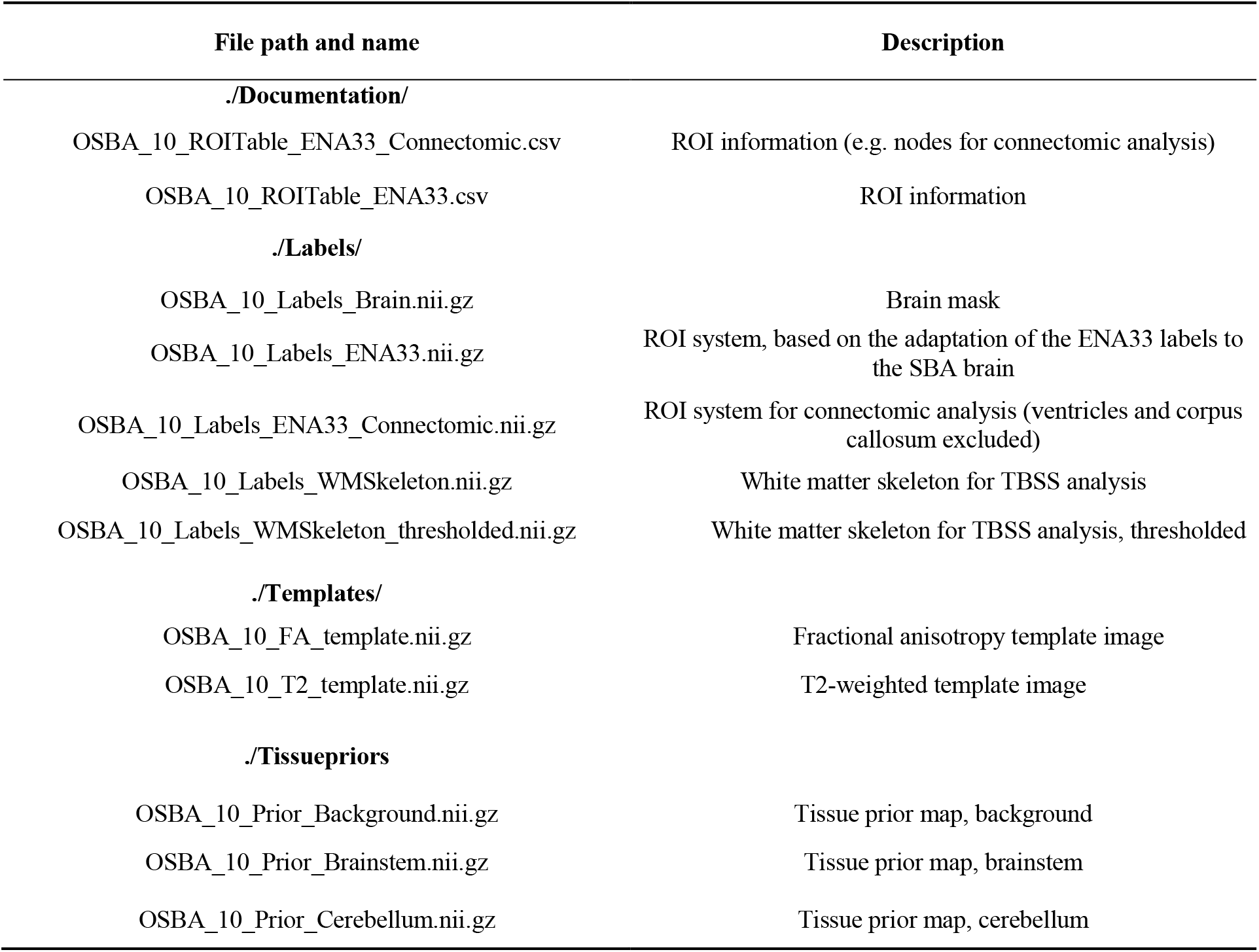
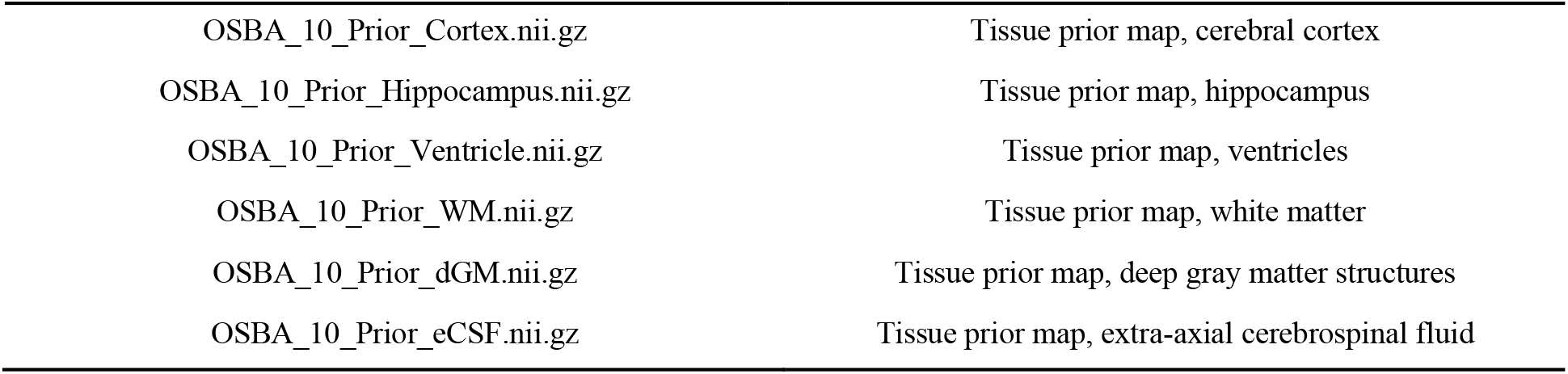
List of files in the OSBA atlas.

### 2.1. Template images and tissue prior maps

All image files are stored in 64-bit floating-point precision NIFTI-1+ images, gzip compressed. The OSBA atlas includes a T2-weighted template image with an image resolution of and size of 117 * 159 * 126 with isotropic voxel dimensions of 0.85 mm, and a corresponding fractional anisotropy template with identical image dimensions. The following tissue priors are included in the atlas: background, extracerebral fluid spaces, cortical gray matter, white matter, ventricles, deep gray matter structures, hippocampus, brainstem, and cerebellum (Figure 1). The tissue prior maps were normalized to take a value between 0 and 1.

**Figure 1.**
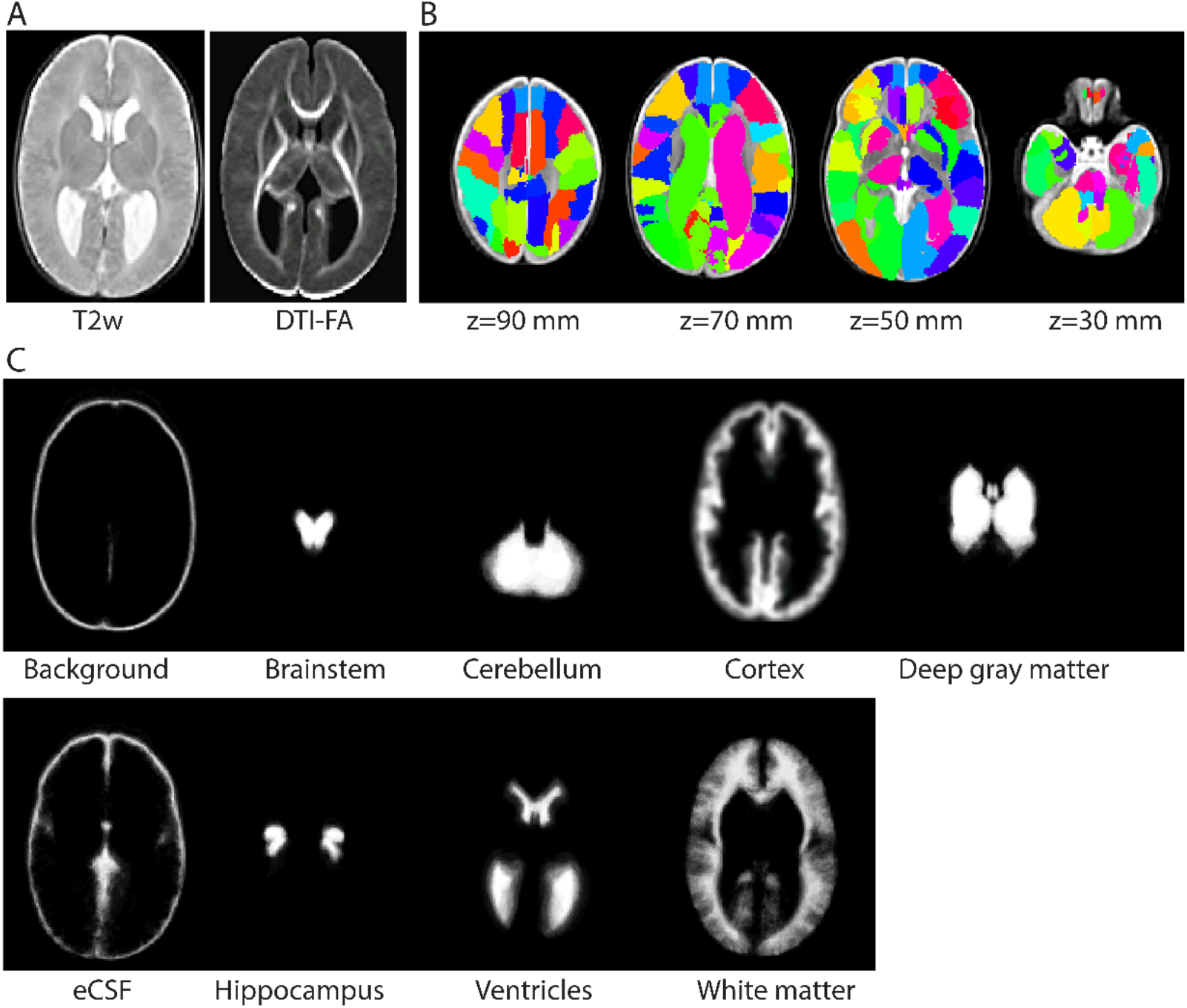
Templates, ROI system and anatomical priors included in the OSBA atlas *Notes*. A: axial cross-section of the T2-weighted and fractional anisotropy anatomical templates, B: ROI label maps at different axial cross-sectional views, C: anatomical prior maps.

### 2.2. Regions of interest (ROI)

Due to the characteristic anatomical configuration of the brain with SBA, the adaptation of ROIs determined from normal subjects is a challenging task. In the SBA atlas, several modifications were made to fit the anatomy of the template images (see 3.5). A recent adaptation of the AAL anatomical ontologies were used by adapting the labels from the publicly available ENA33 atlas to the SBA brain [10]. We include a modified version (‘connectomic’ atlas) of this ROI system in which the labels corresponding to the lateral ventricles and corpus callosum were removed. The label names, spatial coordinates and volumes are detailed in the appendix Table A1 and Table A2.

## 3. Methods

### 3.1. MRI acquisitions

During the neonatal MRI acquisitions, all infants were sedated. Ear protection (earplugs and Minimuffs) was used, oxygen saturation and heart rate were monitored, and all examinations were supervised by a neonatologist or a neonatal nurse. Structural MRI was acquired with fast spin-echo T2-weighted FSE anatomical sequences in axial, coronal, and FRFSE sequence in the sagittal planes on either a 1.5T MRI scanner (GE Signa Discovery MR450) or a 3.0 T MRI scanner using an 8-channel head coil (GE Signa Discovery MR750). 9 infants were scanned on the 1.5T MRI, while 19 were scanned on the 3.0T MRI. The scanning parameters were the following.

T2-weighted MRI was performed in axial, coronal, and sagittal planes for the 1.5T sequence using the following parameters: : 4400-4575 ms (variable), TE: 101-105 ms (variable), flip angle: 160°. Field of view (FOV): 180 mm, acquisition matrix: 320 * 256 (70% sampling); resampled in-plane resolution was 0.35 * 0.35 mm, slice thickness: 2.5 mm with a 0.2 mm slice gap. For the 3.0T sequence, T2-weighted MRI was acquired in the axial plane with TR: 7800-7900 ms (variable), TE: 101-105 ms (variable), flip angle: 111°. FOV: 180 mm, acquisition matrix: 384 * 320 (70% sampling); resampled in-plane resolution was 0.35 * 0.35 mm, slice thickness: 2.5 mm with a 0.2 mm slice gap.

Diffusion tensor imaging (DTI) was acquired in axial plane using a pulsed gradient spin-echo echo-planar imaging (EPI) sequence. 1.5T DTI sequence parameters: TR: 4100-4400 ms (variable), TE: 90-100 ms (variable), acquisition matrix: 128 * 128, field of view: 180 mm, flip angle: 90°. Diffusion weighting was achieved by using 21 diffusion encoding gradient directions at a b-value of 1000 s/mm2 and one b=0 images. Acquisition matrix: 128 * 128; resampled in-plane resolution 0.7 * 0.7 mm, slice thickness: 3 mm, no slice gap. 3.0T DTI sequence parameters: TR: 4100-4400 ms (variable), TE: 100 ms, acquisition matrix: 128 * 128, field of view: 180 mm, flip angle: 90°. Diffusion weighting was achieved by using 35 diffusion encoding gradient directions at a b-value of 700 s/mm2 and four b=0 images. Acquisition matrix: 128 * 128; resampled in-plane resolution 0.7 * 0.7 mm, slice thickness: 3 mm, no slice gap.

### 3.2. Subjects

MRI was performed based on clinical indications. The criteria for undergoing prenatal repair surgery were established based on modified MOMS criteria, as outlined in previous publications authored by our research team [11,12]. The demographics of the first 150 subjects in the clinical cohort study is detailed in the publications of the clinical team [11]. For the construction of the OSBA atlas, we included MRI data of newborns who underwent surgical corrections between 2015 and 2018. The inclusion criteria were: (1) excellent image quality based on visual assessment and (2) lateral ventricle width <15 mm, measured at the trigonum level of the axial T2 MRI. This selection resulted in 28 MRI datasets used for the atlas construction.

The corrected gestational age at the time of the MRI scan was 38.1 ± 1.1 (mean ± SD, range: 35.7 – 40.7), the birth weight of the infants was 2598 ± 445 g (mean ± SD, range: 1400 – 3350 g). The male / female ratio was 16/12. 16 infants had myelomeningocele lesion (57%), while 12 had myeloschisis (43%). The (highest) anatomical levels of the spinal lesion levels were the following (case numbers): L1: 3, L2: 0, L3. 5, L4: 10, L5: 8, S1: 2. Subependymal nodules were reported in 3 infants (10.7%).

### 3.3. MRI processing

For each subject, the super-resolution slice-to-volume reconstruction algorithm SVRTK was applied to the axial, coronal and sagittal T2 images, resulting in a 3D super-resolution reconstructed T2-weighted image (further named as a 3DT2 image) with an isotropic image resolution of 0.4 * 0.4 * 0.4 mm [13]. Next, the 3DT2 images were segmented into anatomical compartments according to the definition of the developing Human Connectome Project (dHCP) structural pipeline [14]. We utilized an in-house network based on the U-Net architecture which was trained on 3DT2 images and ground truth image labels sampled from a population of normally developing neonatal controls and SBA data. Here, the goal was to improve the accuracy of the dHCP pipeline for neonatal brains with SBA. The training data were created in two steps. First, by running the dHCP structural pipeline on the selected normally developing controls from the dHCP dataset release v2 and SBA cases from Zürich. Next, we carried out manual corrections for cases with errors, particularly in the presence of ventricular enlargement. The network was then retrained on the corrected dataset and further used for segmenting the SBA data for the OSBA atlas.

DTI was processed using an in-house script wrapping common image processing libraries. For slice-to-volume reconstruction and to correct for eddy current and head motion-induced geometric distortions, the eddy_cuda in the Functional Magnetic Imaging of the Brain Software Library (FSL) was used [15]. Next, the dtifit command in FSL was utilized for diffusion tensor and scalar maps estimation (such as fractional anisotropy – FA, B0 image and mean diffusivity map), scalars were estimated using weighted least squares regression.

### 3.4. Template construction

The initial spatial reference of the OSBA atlas was the T2-weighted template from the ALBERTS atlas representing the average normally developing neonatal brain at the 38. corrected gestation week [16], matching the mean age at MRI of the subjects in our study.

The data to be included in the OSBA atlas were selected to represent the average ventricular dilation excluding cases with pronounced hydrocephalus (ventricular width measured at the trigonum <15 mm), but also selected to be of good to excellent image quality after visual assessment. First, 28 selected subjects’ 3D T2 images were re-oriented to the 38. week T2-weighted template from the neonatal atlas by using a 6 degrees of freedom registration with the flirt tool in FSL [15]. The neonatal template served only as a spatial reference for initial alignment and defining image dimensions.

Next, an unbiased non-linear template representative of the selected subjects was re-constructed using the Advanced Normalization Tools (ANTs) by running the multimodal template reconstruction BASH script included in the software [17,18] with default settings used for template construction.

The B0 images (intrinsically carrying T2-weighted contrast) from the DTI were transformed to the same subject’s 3DT2 image using a 12 degrees of freedom registration with the flirt tool in FSL. Next, we used the transformations established in the previous template construction steps to match the DTI space to the T2-weighted template’s space. We transformed the FA maps to the T2-weighted template, and the median image across all subjects were estimated to reduce the effects of data outliers and the result image was saved as the FA template. A white matter skeleton was derived from this FA template by using the tbss_skeleton command implemented in FSL [19].

### 3.5. Anatomical parcellation and transformation of common neonatal ROIs to the OSBA atlas

First, probabilistic maps of anatomical tissue priors were created. We saved the binary label maps corresponding to each of the segmentation labels, and then transformed these using the non-linear deformation to match them to the template space. The average map was then divided by its maximum value to normalize it to take values between 0 and 1.

The OSBA atlas includes anatomical ROIs based on the ENA33 neonatal atlas [10]. The ENA33 comprises T2-weighted templates, which were matched to the SBA T2-weighted template. However, in this process, we noted that neither a linear nor local non-linear registration, in which the registration metrics are globally optimized for the whole anatomical image, were able to precisely match the borders of the subcortical tissues between the anatomical template of the ENA33 and the SBA atlas. This was mainly attributed to the enlarged ventricles and the characteristic shape of the lateral ventricles and brainstem. Therefore, we split the ROIs into groups corresponding to the cortex, subcortical gray matter structures, and other structures.

All cortical ROIs were transformed from the ENA33 to the SBA by non-linear registration matching the probability maps corresponding to the gray matter. Subcortical ROIs, due to the adjacency to the dilated ventricles, were first automatically, then manually registered to the SBA template separately in the following three groups: the bilateral caudate nucleus (label 71,72), cingulate ROIs (labels 31-36) and ROIs corresponding to the thalamus, pallidum, putamen, and hippocampus. These registrations were carried out in the Slicer 3D software with a 6 degrees of freedom registration and the registration accuracy was visually controlled [20]. The rest of the ROIs were transformed to the SBA by image-based (global) registrations guided by matching the T2-weighted templates in both atlases. After this procedure, the ROIs were combined into a single label map.

### 3.6. Limitations

The MRI data utilized in the OSBA atlas is restricted to the early neonatal period because MRI scans were clinically indicated during this age range. Consequently, there is no opportunity for a cross-sectional representation of early brain development, as in our clinic, MRI is acquired at one neonatal time point, preferably within 2 to 4 weeks after birth. A further limitation is that the OSBA atlas only contains asymmetric templates and tissue priors. Future works should establish symmetric templates for less biased analysis of left-right asymmetries in the brain with SBA. The OSBA atlas falls short in capturing the variability present in brains affected by SBA. Conditional atlas generation methods have the capacity to represent the diverse anatomical variations observed in SBA. In the context of SBA, this would mean the different grades of ventriculomegaly, Chiari-II grade, and the presence of various forms of corpus callosum abnormalities associated with SBA. Currently, we excluded cases with severe ventriculomegaly due to the difficulties in co-registering them with images of mild or moderate ventriculomegaly. However, future versions of the OSBA atlas might include such representations of suchvariability in SBA. Lastly, the OSBA atlas is limited to two MRI sequences, T2-weighted and diffusion tensor imaging.

## Author Contributions

Conceptualization, A.S. and A.J.; methodology, A.S. and A.J.; software, A.S. and A.J.; validation, A.S., K.P. and A.J.; formal analysis, A.S. and A.J..; investigation, A.S. and A.J.; resources, L.M., B.P., U.M., K.P. and A.J; data curation, S.A., H.J. and A.J.; writing—original draft preparation, A.S., K.P. and A.J.; writing—review and editing, A.S., H.J., L.M., B.P., U.M. and A.J.; visualization, A.J.; supervision, A.J.; project administration, A.J.; funding acquisition, A. J. All authors have read and agreed to the published version of the manuscript.

## Funding

This research was funded by URPP Adaptive Brain Circuits in Development and Learning (AdaBD) project, the Prof. Dr Max Cloetta Foundation, the EMDO Foundation, and the Vontobel Foundation.

## Institutional Review Board Statement

All parents or caregivers gave written informed consent for the further use of their infants’ data in research. The study was conducted in accordance with the Declaration of Helsinki, and approved by the Ethics Committee of the Canton of Zurich for collecting and analyzing data retrospectively (decision numbers: 2016-01019, 2021-01101 and 2022-0115). Informed Consent Statement: Informed consent was obtained from all subjects involved in the study.

## Data Availability Statement

The data presented in this study are available on Zenodo on request from the corresponding author due to restrictions imposed to ensure compliance with research ethics and data privacy regulations.

## Acknowledgments

The authors want to first thank all families who participated in this research. Infrastructure support for this research was provided by the Clinical Trial Center, University Hospital of Zurich.

The authors want to first thank all families who participated in this research. In addition, we thank our contributing study group without whom this research would not be possible. From the University Children’s Hospital this includes Barbara Casanova, Ruth Etter, Patrice Grehten, Domenic Grisch, Cornelia Hagmann, Mirjam Harm, Maya Horst Luethy, Jenny Kienzler, Raimund Kottke, Niklaus Krayenbuehl, Markus A. Landolt, Bea Latal Hajnal; Andreas Meyer-Heim, Theres Moehrlen, Svea Muehlberg, Evelyne Riesen, Brigitte Seliner, Mithula Shellvarajah, Alexandra Wattinger and Noemi Zweifel. From the University Hospital Zurich, our study group consists of Lukas Kandler, Nicole Ochsenbein, Nele Struebing, Max Antonio Thomasius and Ladina Vonzun. Infrastructure support for this research was provided by the Clinical Trial Center, University Hospital of Zurich.

## Conflicts of Interest

The authors declare no conflicts of interest. The funders had no role in the design of the study; in the collection, analyses, or interpretation of data; in the writing of the manuscript; or in the decision to publish the results.

## Appendix

**Table A1.** ROI information of the OSBA atlas.

| Label | Label name | COG in voxel dimensions |  |  | COG in mm dimensions |  |  | Volume in mm<sup>3</sup> |  |
| --- | --- | --- | --- | --- | --- | --- | --- | --- | --- |
|  |  | x | y | z | x | y | z | OSBA | ENA33 |
| 1 | Precentral_L | 85.543 | 86.576 | 88.759 | -21.951 | -9.770 | 37.558 | 11166 | 10257 |
| 2 | Precentral_R | 32.970 | 90.215 | 88.389 | 23.229 | -6.643 | 37.240 | 11029 | 10403 |
| 3 | Frontal_Sup_L | 72.947 | 115.517 | 82.997 | -11.126 | 15.101 | 32.607 | 13462 | 11547 |
| 4 | Frontal_Sup_R | 46.372 | 119.390 | 79.514 | 11.712 | 18.429 | 29.613 | 11290 | 10134 |
| 5 | Frontal_Sup_Orb_L | 68.917 | 122.058 | 39.058 | -7.663 | 20.721 | -5.154 | 2134 | 1824 |
| 6 | Frontal_Sup_Orb_R | 50.279 | 120.012 | 38.082 | 8.354 | 18.963 | -5.992 | 1983 | 1668 |
| 7 | Frontal_Mid_L | 83.561 | 113.941 | 75.008 | -20.248 | 13.746 | 25.741 | 15545 | 13495 |
| 8 | Frontal_Mid_R | 35.854 | 115.685 | 75.964 | 20.750 | 15.245 | 26.563 | 16746 | 15345 |
| 9 | Frontal_Mid_Orb_L | 76.690 | 128.197 | 42.521 | -14.343 | 25.998 | -2.177 | 3193 | 2635 |
| 10 | Frontal_Mid_Orb_R | 41.864 | 127.568 | 42.813 | 15.586 | 25.457 | -1.926 | 3237 | 2902 |
| 11 | Frontal_Inf_Oper_L | 90.597 | 101.307 | 63.473 | -26.294 | 2.889 | 15.828 | 3671 | 3332 |
| 12 | Frontal_Inf_Oper_R | 27.712 | 103.866 | 62.958 | 27.748 | 5.088 | 15.386 | 3695 | 3367 |
| 13 | Frontal_Inf_Tri_L | 89.779 | 113.174 | 57.294 | -25.591 | 13.087 | 10.518 | 3793 | 3392 |
| 14 | Frontal_Inf_Tri_R | 28.135 | 115.343 | 57.695 | 27.384 | 14.951 | 10.863 | 4082 | 3727 |
| 15 | Frontal_Inf_Orb_L | 82.036 | 111.898 | 42.482 | -18.937 | 11.991 | -2.210 | 5958 | 6071 |
| 16 | Frontal_Inf_Orb_R | 36.759 | 113.415 | 42.396 | 19.973 | 13.294 | -2.285 | 4928 | 4887 |
| 17 | Rolandic_Oper_L | 93.084 | 81.361 | 66.223 | -28.432 | -14.253 | 18.192 | 7764 | 6556 |
| 18 | Rolandic_Oper_R | 25.235 | 84.754 | 67.886 | 29.876 | -11.337 | 19.621 | 6111 | 4549 |
| 19 | Supp_Motor_Area_L | 64.366 | 94.133 | 96.885 | -3.752 | -3.276 | 44.542 | 7037 | 7196 |
| 20 | Supp_Motor_Area_R | 53.794 | 95.604 | 96.514 | 5.334 | -2.012 | 44.223 | 5880 | 6663 |
| 21 | Olfactory_L | 65.126 | 100.927 | 41.292 | -4.405 | 2.562 | -3.234 | 1665 | 1543 |
| 22 | Olfactory_R | 53.974 | 101.550 | 41.781 | 5.178 | 3.097 | -2.813 | 1818 | 1630 |
| 23 | Frontal_Sup_Medial_L | 64.617 | 127.752 | 71.887 | -3.968 | 25.615 | 23.059 | 7932 | 7133 |
| 24 | Frontal_Sup_Medial_R | 54.444 | 127.603 | 71.539 | 4.775 | 25.487 | 22.760 | 8383 | 8390 |
| 25 | Frontal_Med_Orb_L | 63.617 | 122.037 | 47.188 | -3.108 | 20.703 | 1.834 | 3836 | 1990 |
| 26 | Frontal_Med_Orb_R | 54.542 | 124.243 | 45.045 | 4.690 | 22.599 | -0.008 | 3262 | 1973 |
| 27 | Rectus_L | 64.399 | 117.876 | 36.548 | -3.781 | 17.128 | -7.311 | 1603 | 1354 |
| 28 | Rectus_R | 55.203 | 118.714 | 36.443 | 4.122 | 17.848 | -7.400 | 1457 | 1301 |
| 29 | Insula_L | 82.997 | 94.258 | 53.186 | -19.763 | -3.169 | 6.988 | 3757 | 4901 |
| 30 | Insula_R | 31.986 | 95.366 | 53.804 | 24.074 | -2.217 | 7.519 | 2026 | 4501 |
| 31 | Cingulum_Ant_L | 62.574 | 111.255 | 69.403 | -2.212 | 11.438 | 20.924 | 2393 | 3089 |
| 32 | Cingulum_Ant_R | 55.256 | 113.158 | 63.890 | 4.077 | 13.073 | 16.186 | 3036 | 3003 |
| 33 | Cingulum_Mid_L | 62.440 | 71.317 | 84.300 | -2.097 | -22.884 | 33.727 | 3654 | 4472 |
| 34 | Cingulum_Mid_R | 52.928 | 71.652 | 84.377 | 6.077 | -22.596 | 33.793 | 3651 | 4816 |
| 35 | Cingulum_Post_L | 61.136 | 56.598 | 72.407 | -0.976 | -35.533 | 23.506 | 199 | 454 |
| 36 | Cingulum_Post_R | 55.820 | 57.753 | 74.127 | 3.592 | -34.541 | 24.984 | 267 | 565 |
| 37 | Hippocampus_L | 77.737 | 77.931 | 40.158 | -15.242 | -17.200 | -4.208 | 1732 | 2658 |
| 38 | Hippocampus_R | 40.475 | 80.128 | 41.675 | 16.780 | -15.312 | -2.904 | 2073 | 2793 |
| 39 | ParaHippocampal_L | 74.656 | 80.363 | 34.580 | -12.595 | -15.110 | -9.002 | 2407 | 3193 |
| 40 | ParaHippocampal_R | 43.663 | 82.537 | 34.942 | 14.039 | -13.241 | -8.691 | 2313 | 3039 |
| 41 | Amygdala_L | 76.662 | 89.686 | 36.916 | -14.319 | -7.098 | -6.994 | 509 | 821 |
| 42 | Amygdala_R | 39.625 | 92.468 | 38.109 | 17.510 | -4.707 | -5.969 | 523 | 963 |
| 43 | Calcarine_L | 61.944 | 27.103 | 62.783 | -1.671 | -60.880 | 15.235 | 5354 | 5546 |
| 44 | Calcarine_R | 50.685 | 32.047 | 64.413 | 8.005 | -56.631 | 16.636 | 5457 | 5189 |
| 45 | Cuneus_L | 64.107 | 29.395 | 78.938 | -3.530 | -58.911 | 29.119 | 5099 | 4541 |
| 46 | Cuneus_R | 49.818 | 33.323 | 79.395 | 8.751 | -55.535 | 29.511 | 5047 | 4394 |
| 47 | Lingual_L | 65.725 | 31.759 | 48.574 | -4.920 | -56.879 | 3.024 | 9004 | 9341 |
| 48 | Lingual_R | 48.215 | 33.355 | 49.769 | 10.128 | -55.507 | 4.051 | 10212 | 9726 |
| 49 | Occipital_Sup_L | 73.440 | 30.657 | 80.139 | -11.550 | -57.826 | 30.151 | 3126 | 2780 |
| 50 | Occipital_Sup_R | 41.154 | 28.882 | 78.842 | 16.196 | -59.352 | 29.036 | 4698 | 4221 |
| 51 | Occipital_Mid_L | 79.339 | 26.437 | 69.128 | -16.619 | -61.453 | 20.688 | 7811 | 7164 |
| 52 | Occipital_Mid_R | 31.690 | 33.049 | 69.240 | 24.329 | -55.770 | 20.785 | 8067 | 7933 |
| 53 | Occipital_Inf_L | 83.249 | 26.610 | 50.274 | -19.980 | -61.304 | 4.486 | 4258 | 4156 |
| 54 | Occipital_Inf_R | 30.633 | 30.554 | 51.783 | 25.237 | -57.914 | 5.783 | 7272 | 6727 |
| 55 | Fusiform_L | 80.795 | 57.900 | 37.179 | -17.871 | -34.414 | -6.768 | 6442 | 6906 |
| 56 | Fusiform_R | 35.314 | 58.065 | 39.493 | 21.214 | -34.272 | -4.780 | 5727 | 6407 |
| 57 | Postcentral_L | 86.510 | 73.361 | 91.951 | -22.782 | -21.127 | 40.302 | 9347 | 8252 |
| 58 | Postcentral_R | 31.164 | 76.934 | 92.571 | 24.781 | -18.057 | 40.834 | 9599 | 8597 |
| 59 | Parietal_Sup_L | 76.020 | 49.592 | 97.510 | -13.767 | -41.554 | 45.079 | 7570 | 6655 |
| 60 | Parietal_Sup_R | 42.249 | 51.269 | 99.056 | 15.255 | -40.112 | 46.407 | 6198 | 5583 |
| 61 | Parietal_Inf_L | 87.763 | 55.315 | 88.647 | -23.859 | -36.636 | 37.462 | 2966 | 2872 |
| 62 | Parietal_Inf_R | 30.081 | 59.050 | 88.504 | 25.712 | -33.426 | 37.339 | 8400 | 7008 |
| 63 | SupraMarginal_L | 95.107 | 62.319 | 79.087 | -30.170 | -30.617 | 29.246 | 5555 | 4723 |
| 64 | SupraMarginal_R | 19.688 | 65.960 | 79.010 | 34.643 | -27.487 | 29.180 | 4253 | 3580 |
| 65 | Angular_L | 87.365 | 42.717 | 81.493 | -23.516 | -47.462 | 31.314 | 4229 | 4237 |
| 66 | Angular_R | 26.293 | 45.882 | 79.961 | 28.967 | -44.742 | 29.998 | 2794 | 2729 |
| 67 | Precuneus_L | 63.791 | 47.390 | 87.647 | -3.258 | -43.446 | 36.603 | 9210 | 8884 |
| 68 | Precuneus_R | 52.067 | 48.190 | 87.674 | 6.817 | -42.759 | 36.626 | 8976 | 8573 |
| 69 | Paracentral_Lobule_L | 63.869 | 67.299 | 102.016 | -3.325 | -26.337 | 48.951 | 2960 | 2733 |
| 70 | Paracentral_Lobule_R | 52.008 | 71.244 | 102.542 | 6.868 | -22.946 | 49.403 | 4133 | 3867 |
| 71 | Caudate_L | 69.778 | 100.837 | 55.905 | -8.403 | 2.485 | 9.324 | 693 | 1362 |
| 72 | Caudate_R | 48.189 | 103.485 | 55.091 | 10.151 | 4.761 | 8.625 | 870 | 1449 |
| 73 | Putamen_L | 77.168 | 95.553 | 52.274 | -14.754 | -2.056 | 6.205 | 1782 | 1995 |
| 74 | Putamen_R | 37.948 | 98.198 | 53.344 | 18.951 | 0.217 | 7.124 | 2077 | 2334 |
| 75 | Pallidum_L | 72.988 | 94.696 | 50.770 | -11.162 | -2.792 | 4.912 | 1652 | 1801 |
| 76 | Pallidum_R | 43.154 | 96.604 | 50.989 | 14.477 | -1.153 | 5.100 | 1712 | 1910 |
| 77 | Thalamus_L | 69.029 | 77.262 | 57.078 | -7.759 | -17.775 | 10.333 | 5915 | 4967 |
| 78 | Thalamus_R | 47.634 | 79.562 | 57.623 | 10.627 | -15.798 | 10.801 | 5825 | 5095 |
| 79 | Heschl_L | 89.732 | 74.852 | 60.298 | -25.551 | -19.846 | 13.100 | 1293 | 1263 |
| 80 | Heschl_R | 28.961 | 79.100 | 61.322 | 26.674 | -16.196 | 13.980 | 1214 | 1134 |
| 81 | Temporal_Sup_L | 95.094 | 68.860 | 58.463 | -30.159 | -24.995 | 11.522 | 8851 | 7889 |
| 82 | Temporal_Sup_R | 22.367 | 76.721 | 58.030 | 32.341 | -18.239 | 11.151 | 9445 | 8941 |
| 83 | Temporal_Pole_Sup_L | 89.355 | 95.144 | 37.500 | -25.227 | -2.408 | -6.492 | 3254 | 3193 |
| 84 | Temporal_Pole_Sup_R | 29.844 | 98.693 | 37.927 | 25.915 | 0.642 | -6.125 | 2600 | 3424 |
| 85 | Temporal_Mid_L | 93.988 | 56.891 | 54.219 | -29.209 | -35.282 | 7.876 | 9974 | 9265 |
| 86 | Temporal_Mid_R | 22.072 | 66.520 | 52.941 | 32.595 | -27.006 | 6.778 | 11112 | 10498 |
| 87 | Temporal_Pole_Mid_L | 87.198 | 94.812 | 26.667 | -23.373 | -2.693 | -15.802 | 1310 | 1698 |
| 88 | Temporal_Pole_Mid_R | 32.747 | 97.307 | 25.565 | 23.421 | -0.548 | -16.749 | 774 | 1524 |
| 89 | Temporal_Inf_L | 91.913 | 61.356 | 39.051 | -27.425 | -31.444 | -5.159 | 10160 | 9878 |
| 90 | Temporal_Inf_R | 25.423 | 68.582 | 36.859 | 29.714 | -25.234 | -7.043 | 8351 | 7681 |
| 91 | CorpusCallosum | 57.581 | 85.064 | 73.483 | 2.079 | -11.070 | 24.431 | 4330 | 3562 |
| 92 | Lateral_Ventricle_L | 72.414 | 68.197 | 65.837 | -10.668 | -25.565 | 17.860 | 27034 | 3228 |
| 93 | Lateral_Ventricle_R | 43.415 | 70.915 | 66.878 | 14.252 | -23.230 | 18.755 | 22297 | 3361 |
| 94 | Midbrain_L | 63.015 | 72.828 | 42.660 | -2.591 | -21.585 | -2.058 | 1854 | 2341 |
| 95 | Midbrain_R | 54.308 | 73.016 | 42.968 | 4.891 | -21.423 | -1.793 | 1707 | 2331 |
| 96 | Pons_L | 62.511 | 72.080 | 29.032 | -2.158 | -22.228 | -13.770 | 1007 | 1067 |
| 97 | Pons_R | 54.973 | 71.696 | 28.977 | 4.320 | -22.558 | -13.817 | 1212 | 1347 |
| 98 | Medulla_L | 63.423 | 61.522 | 23.212 | -2.942 | -31.301 | -18.771 | 2113 | 2582 |
| 99 | Medulla_R | 55.003 | 61.331 | 22.581 | 4.294 | -31.466 | -19.313 | 2369 | 3044 |
| 100 | Cerebellum_L | 71.631 | 48.908 | 28.988 | -9.995 | -42.142 | -13.807 | 11410 | 13721 |
| 101 | Cerebellum_R | 44.934 | 49.781 | 29.499 | 12.947 | -41.392 | -13.368 | 11423 | 13704 |
| 102 | Vermis_Ant_L | 59.958 | 53.432 | 39.882 | 0.036 | -38.254 | -4.445 | 542 | 935 |
| 103 | Vermis_Ant_R | 56.278 | 53.478 | 40.116 | 3.199 | -38.214 | -4.244 | 508 | 914 |
| 104 | Vermis_Post_L | 63.180 | 47.630 | 31.171 | -2.733 | -43.240 | -11.931 | 1680 | 2353 |
| 105 | Vermis_Post_R | 54.301 | 47.333 | 31.114 | 4.897 | -43.495 | -11.980 | 1470 | 2297 |
| 106 | Vermis_Cent_L | 62.153 | 51.094 | 24.805 | -1.850 | -40.263 | -17.402 | 615 | 880 |
| 107 | Vermis_Cent_R | 54.236 | 52.135 | 24.390 | 4.953 | -39.369 | -17.759 | 728 | 1053 |

Notes. COG = center of gravity.

**Table A2.** ROI information of the connectomic OSBA atlas.

| Label | Label name | COG in voxel dimensions |  |  | COG in mm dimensions |  |  | Volume in mm<sup>3</sup> |  |
| --- | --- | --- | --- | --- | --- | --- | --- | --- | --- |
|  |  | x | y | z | x | y | z | OSBA | ENA33 |
| 1 | Precentral_L | 85.543 | 86.576 | 88.759 | -21.951 | -9.770 | 37.558 | 11166 | 10257 |
| 2 | Precentral_R | 32.970 | 90.215 | 88.389 | 23.229 | -6.643 | 37.240 | 11029 | 10403 |
| 3 | Frontal_Sup_L | 72.947 | 115.517 | 82.997 | -11.126 | 15.101 | 32.607 | 13462 | 11547 |
| 4 | Frontal_Sup_R | 46.372 | 119.390 | 79.514 | 11.712 | 18.429 | 29.613 | 11290 | 10134 |
| 5 | Frontal_Sup_Orb_L | 68.917 | 122.058 | 39.058 | -7.663 | 20.721 | -5.154 | 2134 | 1824 |
| 6 | Frontal_Sup_Orb_R | 50.279 | 120.012 | 38.082 | 8.354 | 18.963 | -5.992 | 1983 | 1668 |
| 7 | Frontal_Mid_L | 83.561 | 113.941 | 75.008 | -20.248 | 13.746 | 25.741 | 15545 | 13495 |
| 8 | Frontal_Mid_R | 35.854 | 115.685 | 75.964 | 20.750 | 15.245 | 26.563 | 16746 | 15345 |
| 9 | Frontal_Mid_Orb_L | 76.690 | 128.197 | 42.521 | -14.343 | 25.998 | -2.177 | 3193 | 2635 |
| 10 | Frontal_Mid_Orb_R | 41.864 | 127.568 | 42.813 | 15.586 | 25.457 | -1.926 | 3237 | 2902 |
| 11 | Frontal_Inf_Oper_L | 90.597 | 101.307 | 63.473 | -26.294 | 2.889 | 15.828 | 3671 | 3332 |
| 12 | Frontal_Inf_Oper_R | 27.712 | 103.866 | 62.958 | 27.748 | 5.088 | 15.386 | 3695 | 3367 |
| 13 | Frontal_Inf_Tri_L | 89.779 | 113.174 | 57.294 | -25.591 | 13.087 | 10.518 | 3793 | 3392 |
| 14 | Frontal_Inf_Tri_R | 28.135 | 115.343 | 57.695 | 27.384 | 14.951 | 10.863 | 4082 | 3727 |
| 15 | Frontal_Inf_Orb_L | 82.036 | 111.898 | 42.482 | -18.937 | 11.991 | -2.210 | 5958 | 6071 |
| 16 | Frontal_Inf_Orb_R | 36.759 | 113.415 | 42.396 | 19.973 | 13.294 | -2.285 | 4928 | 4887 |
| 17 | Rolandic_Oper_L | 93.084 | 81.361 | 66.223 | -28.432 | -14.253 | 18.192 | 7764 | 6556 |
| 18 | Rolandic_Oper_R | 25.235 | 84.754 | 67.886 | 29.876 | -11.337 | 19.621 | 6111 | 4549 |
| 19 | Supp_Motor_Area_L | 64.366 | 94.133 | 96.885 | -3.752 | -3.276 | 44.542 | 7037 | 7196 |
| 20 | Supp_Motor_Area_R | 53.794 | 95.604 | 96.514 | 5.334 | -2.012 | 44.223 | 5880 | 6663 |
| 21 | Olfactory_L | 65.126 | 100.927 | 41.292 | -4.405 | 2.562 | -3.234 | 1665 | 1543 |
| 22 | Olfactory_R | 53.974 | 101.550 | 41.781 | 5.178 | 3.097 | -2.813 | 1818 | 1630 |
| 23 | Frontal_Sup_Medial_L | 64.617 | 127.752 | 71.887 | -3.968 | 25.615 | 23.059 | 7932 | 7133 |
| 24 | Frontal_Sup_Medial_R | 54.444 | 127.603 | 71.539 | 4.775 | 25.487 | 22.760 | 8383 | 8390 |
| 25 | Frontal_Med_Orb_L | 63.617 | 122.037 | 47.188 | -3.108 | 20.703 | 1.834 | 3836 | 1990 |
| 26 | Frontal_Med_Orb_R | 54.542 | 124.243 | 45.045 | 4.690 | 22.599 | -0.008 | 3262 | 1973 |
| 27 | Rectus_L | 64.399 | 117.876 | 36.548 | -3.781 | 17.128 | -7.311 | 1603 | 1354 |
| 28 | Rectus_R | 55.203 | 118.714 | 36.443 | 4.122 | 17.848 | -7.400 | 1457 | 1301 |
| 29 | Insula_L | 82.997 | 94.258 | 53.186 | -19.763 | -3.169 | 6.988 | 3757 | 4901 |
| 30 | Insula_R | 31.986 | 95.366 | 53.804 | 24.074 | -2.217 | 7.519 | 2026 | 4501 |
| 31 | Cingulum_Ant_L | 62.574 | 111.255 | 69.403 | -2.212 | 11.438 | 20.924 | 2393 | 3089 |
| 32 | Cingulum_Ant_R | 55.256 | 113.158 | 63.890 | 4.077 | 13.073 | 16.186 | 3036 | 3003 |
| 33 | Cingulum_Mid_L | 62.440 | 71.317 | 84.300 | -2.097 | -22.884 | 33.727 | 3654 | 4472 |
| 34 | Cingulum_Mid_R | 52.928 | 71.652 | 84.377 | 6.077 | -22.596 | 33.793 | 3651 | 4816 |
| 35 | Cingulum_Post_L | 61.136 | 56.598 | 72.407 | -0.976 | -35.533 | 23.506 | 199 | 454 |
| 36 | Cingulum_Post_R | 55.820 | 57.753 | 74.127 | 3.592 | -34.541 | 24.984 | 267 | 565 |
| 37 | Hippocampus_L | 77.737 | 77.931 | 40.158 | -15.242 | -17.200 | -4.208 | 1732 | 2658 |
| 38 | Hippocampus_R | 40.475 | 80.128 | 41.675 | 16.780 | -15.312 | -2.904 | 2073 | 2793 |
| 39 | ParaHippocampal_L | 74.656 | 80.363 | 34.580 | -12.595 | -15.110 | -9.002 | 2407 | 3193 |
| 40 | ParaHippocampal_R | 43.663 | 82.537 | 34.942 | 14.039 | -13.241 | -8.691 | 2313 | 3039 |
| 41 | Amygdala_L | 76.662 | 89.686 | 36.916 | -14.319 | -7.098 | -6.994 | 509 | 821 |
| 42 | Amygdala_R | 39.625 | 92.468 | 38.109 | 17.510 | -4.707 | -5.969 | 523 | 963 |
| 43 | Calcarine_L | 61.944 | 27.103 | 62.783 | -1.671 | -60.880 | 15.235 | 5354 | 5546 |
| 44 | Calcarine_R | 50.685 | 32.047 | 64.413 | 8.005 | -56.631 | 16.636 | 5457 | 5189 |
| 45 | Cuneus_L | 64.107 | 29.395 | 78.938 | -3.530 | -58.911 | 29.119 | 5099 | 4541 |
| 46 | Cuneus_R | 49.818 | 33.323 | 79.395 | 8.751 | -55.535 | 29.511 | 5047 | 4394 |
| 47 | Lingual_L | 65.725 | 31.759 | 48.574 | -4.920 | -56.879 | 3.024 | 9004 | 9341 |
| 48 | Lingual_R | 48.215 | 33.355 | 49.769 | 10.128 | -55.507 | 4.051 | 10212 | 9726 |
| 49 | Occipital_Sup_L | 73.440 | 30.657 | 80.139 | -11.550 | -57.826 | 30.151 | 3126 | 2780 |
| 50 | Occipital_Sup_R | 41.154 | 28.882 | 78.842 | 16.196 | -59.352 | 29.036 | 4698 | 4221 |
| 51 | Occipital_Mid_L | 79.339 | 26.437 | 69.128 | -16.619 | -61.453 | 20.688 | 7811 | 7164 |
| 52 | Occipital_Mid_R | 31.690 | 33.049 | 69.240 | 24.329 | -55.770 | 20.785 | 8067 | 7933 |
| 53 | Occipital_Inf_L | 83.249 | 26.610 | 50.274 | -19.980 | -61.304 | 4.486 | 4258 | 4156 |
| 54 | Occipital_Inf_R | 30.633 | 30.554 | 51.783 | 25.237 | -57.914 | 5.783 | 7272 | 6727 |
| 55 | Fusiform_L | 80.795 | 57.900 | 37.179 | -17.871 | -34.414 | -6.768 | 6442 | 6906 |
| 56 | Fusiform_R | 35.314 | 58.065 | 39.493 | 21.214 | -34.272 | -4.780 | 5727 | 6407 |
| 57 | Postcentral_L | 86.510 | 73.361 | 91.951 | -22.782 | -21.127 | 40.302 | 9347 | 8252 |
| 58 | Postcentral_R | 31.164 | 76.934 | 92.571 | 24.781 | -18.057 | 40.834 | 9599 | 8597 |
| 59 | Parietal_Sup_L | 76.020 | 49.592 | 97.510 | -13.767 | -41.554 | 45.079 | 7570 | 6655 |
| 60 | Parietal_Sup_R | 42.249 | 51.269 | 99.056 | 15.255 | -40.112 | 46.407 | 6198 | 5583 |
| 61 | Parietal_Inf_L | 87.763 | 55.315 | 88.647 | -23.859 | -36.636 | 37.462 | 2966 | 2872 |
| 62 | Parietal_Inf_R | 30.081 | 59.050 | 88.504 | 25.712 | -33.426 | 37.339 | 8400 | 7008 |
| 63 | SupraMarginal_L | 95.107 | 62.319 | 79.087 | -30.170 | -30.617 | 29.246 | 5555 | 4723 |
| 64 | SupraMarginal_R | 19.688 | 65.960 | 79.010 | 34.643 | -27.487 | 29.180 | 4253 | 3580 |
| 65 | Angular_L | 87.365 | 42.717 | 81.493 | -23.516 | -47.462 | 31.314 | 4229 | 4237 |
| 66 | Angular_R | 26.293 | 45.882 | 79.961 | 28.967 | -44.742 | 29.998 | 2794 | 2729 |
| 67 | Precuneus_L | 63.791 | 47.390 | 87.647 | -3.258 | -43.446 | 36.603 | 9210 | 8884 |
| 68 | Precuneus_R | 52.067 | 48.190 | 87.674 | 6.817 | -42.759 | 36.626 | 8976 | 8573 |
| 69 | Paracentral_Lobule_L | 63.869 | 67.299 | 102.016 | -3.325 | -26.337 | 48.951 | 2960 | 2733 |
| 70 | Paracentral_Lobule_R | 52.008 | 71.244 | 102.542 | 6.868 | -22.946 | 49.403 | 4133 | 3867 |
| 71 | Caudate_L | 69.778 | 100.837 | 55.905 | -8.403 | 2.485 | 9.324 | 693 | 1362 |
| 72 | Caudate_R | 48.189 | 103.485 | 55.091 | 10.151 | 4.761 | 8.625 | 870 | 1449 |
| 73 | Putamen_L | 77.168 | 95.553 | 52.274 | -14.754 | -2.056 | 6.205 | 1782 | 1995 |
| 74 | Putamen_R | 37.948 | 98.198 | 53.344 | 18.951 | 0.217 | 7.124 | 2077 | 2334 |
| 75 | Pallidum_L | 72.988 | 94.696 | 50.770 | -11.162 | -2.792 | 4.912 | 1652 | 1801 |
| 76 | Pallidum_R | 43.154 | 96.604 | 50.989 | 14.477 | -1.153 | 5.100 | 1712 | 1910 |
| 77 | Thalamus_L | 69.029 | 77.262 | 57.078 | -7.759 | -17.775 | 10.333 | 5915 | 4967 |
| 78 | Thalamus_R | 47.634 | 79.562 | 57.623 | 10.627 | -15.798 | 10.801 | 5825 | 5095 |
| 79 | Heschl_L | 89.732 | 74.852 | 60.298 | -25.551 | -19.846 | 13.100 | 1293 | 1263 |
| 80 | Heschl_R | 28.961 | 79.100 | 61.322 | 26.674 | -16.196 | 13.980 | 1214 | 1134 |
| 81 | Temporal_Sup_L | 95.094 | 68.860 | 58.463 | -30.159 | -24.995 | 11.522 | 8851 | 7889 |
| 82 | Temporal_Sup_R | 22.367 | 76.721 | 58.030 | 32.341 | -18.239 | 11.151 | 9445 | 8941 |
| 83 | Temporal_Pole_Sup_L | 89.355 | 95.144 | 37.500 | -25.227 | -2.408 | -6.492 | 3254 | 3193 |
| 84 | Temporal_Pole_Sup_R | 29.844 | 98.693 | 37.927 | 25.915 | 0.642 | -6.125 | 2600 | 3424 |
| 85 | Temporal_Mid_L | 93.988 | 56.891 | 54.219 | -29.209 | -35.282 | 7.876 | 9974 | 9265 |
| 86 | Temporal_Mid_R | 22.072 | 66.520 | 52.941 | 32.595 | -27.006 | 6.778 | 11112 | 10498 |
| 87 | Temporal_Pole_Mid_L | 87.198 | 94.812 | 26.667 | -23.373 | -2.693 | -15.802 | 1310 | 1698 |
| 88 | Temporal_Pole_Mid_R | 32.747 | 97.307 | 25.565 | 23.421 | -0.548 | -16.749 | 774 | 1524 |
| 89 | Temporal_Inf_L | 91.913 | 61.356 | 39.051 | -27.425 | -31.444 | -5.159 | 10160 | 9878 |
| 90 | Temporal_Inf_R | 25.423 | 68.582 | 36.859 | 29.714 | -25.234 | -7.043 | 8351 | 7681 |
| 91 | Midbrain_L | 63.015 | 72.828 | 42.660 | -2.591 | -21.585 | -2.058 | 1854 | 2341 |
| 92 | Midbrain_R | 54.308 | 73.016 | 42.968 | 4.891 | -21.423 | -1.793 | 1707 | 2331 |
| 93 | Pons_L | 62.511 | 72.080 | 29.032 | -2.158 | -22.228 | -13.770 | 1007 | 1067 |
| 94 | Pons_R | 54.973 | 71.696 | 28.977 | 4.320 | -22.558 | -13.817 | 1212 | 1347 |
| 95 | Medulla_L | 63.423 | 61.522 | 23.212 | -2.942 | -31.301 | -18.771 | 2113 | 2582 |
| 96 | Medulla_R | 55.003 | 61.331 | 22.581 | 4.294 | -31.466 | -19.313 | 2369 | 3044 |
| 97 | Cerebellum_L | 71.631 | 48.908 | 28.988 | -9.995 | -42.142 | -13.807 | 11410 | 13721 |
| 98 | Cerebellum_R | 44.934 | 49.781 | 29.499 | 12.947 | -41.392 | -13.368 | 11423 | 13704 |
| 99 | Vermis_Ant_L | 59.958 | 53.432 | 39.882 | 0.036 | -38.254 | -4.445 | 542 | 935 |
| 100 | Vermis_Ant_R | 56.278 | 53.478 | 40.116 | 3.199 | -38.214 | -4.244 | 508 | 914 |
| 101 | Vermis_Post_L | 63.180 | 47.630 | 31.171 | -2.733 | -43.240 | -11.931 | 1680 | 2353 |
| 102 | Vermis_Post_R | 54.301 | 47.333 | 31.114 | 4.897 | -43.495 | -11.980 | 1470 | 2297 |
| 103 | Vermis_Cent_L | 62.153 | 51.094 | 24.805 | -1.850 | -40.263 | -17.402 | 615 | 880 |
| 104 | Vermis_Cent_R | 54.236 | 52.135 | 24.390 | 4.953 | -39.369 | -17.759 | 728 | 1053 |

Notes. COG = cent of gravity.

## Notes

https://doi.org/10.5281/zenodo.11469304

### Competing Interest Statement

The authors have declared no competing interest.

## References

1. Oishi, K.; Chang, L.; Huang, H. Baby Brain Atlases. Neuroimage 2019, 185, 865–880, doi:10.1016/j.neuroimage.2018.04.003.

2. Fidon, L.; Viola, E.; Mufti, N.; David, A.L.; Melbourne, A.; Demaerel, P.; Ourselin, S.; Vercauteren, T.; Deprest, J.; Aertsen, M. A Spatio-Temporal Atlas of the Developing Fetal Brain with Spina Bifida Aperta. Open Research Europe 2022, 1, 123, doi:10.12688/openreseurope.13914.2.

3. Evans, A.C.; Janke, A.L.; Collins, D.L.; Baillet, S. Brain Templates and Atlases. Neuroimage 2012, 62, 911–922, doi:10.1016/j.neuroimage.2012.01.024.

4. Ciceri, T.; Casartelli, L.; Montano, F.; Conte, S.; Squarcina, L.; Bertoldo, A.; Agarwal, N.; Brambilla, P.; Peruzzo, D. Fetal Brain MRI Atlases and Datasets: A Review. Neuroimage 2024, 292, 120603, doi:10.1016/j.neuroimage.2024.120603.

5. Gholipour, A.; Rollins, C.K.; Velasco-Annis, C.; Ouaalam, A.; Akhondi-Asl, A.; Afacan, O.; Ortinau, C.M.; Clancy, S.; Limperopoulos, C.; Yang, E.; et al. A Normative Spatiotemporal MRI Atlas of the Fetal Brain for Automatic Segmentation and Analysis of Early Brain Growth. Sci Rep 2017, 7, 476, doi:10.1038/s41598-017-00525-w.

6. Khoshnood, B.; Loane, M.; Walle, H. de; Arriola, L.; Addor, M.-C.; Barisic, I.; Beres, J.; Bianchi, F.; Dias, C.; Draper, E.; et al. Long Term Trends in Prevalence of Neural Tube Defects in Europe: Population Based Study. BMJ 2015, h5949, doi:10.1136/bmj.h5949.

7. Schneider, J.; Mohr, N.; Aliatakis, N.; Seidel, U.; John, R.; Promnitz, G.; Spors, B.; Kaindl, A.M. Brain Malformations and Cognitive Performance in Spina Bifida. Dev Med Child Neurol 2021, 63, 295–302, doi:10.1111/dmcn.14717.

8. Mufti, N.; Sacco, A.; Aertsen, M.; Ushakov, F.; Ourselin, S.; Thomson, D.; Deprest, J.; Melbourne, A.; David, A.L. What Brain Abnormalities Can Magnetic Resonance Imaging Detect in Foetal and Early Neonatal Spina Bifida: A Systematic Review. Neuroradiology 2022, 64, 233–245, doi:10.1007/s00234-021-02853-1.

9. Shrot, S.; Soares, B.P.; Whitehead, M.T. Cerebral Diffusivity Changes in Fetuses with Chiari II Malformation. Fetal Diagn Ther 2019, 45, 268–274, doi:10.1159/000490102.

10. Blesa, M.; Serag, A.; Wilkinson, A.G.; Anblagan, D.; Telford, E.J.; Pataky, R.; Sparrow, S.A.; Macnaught, G.; Semple, S.I.; Bastin, M.E.; et al. Parcellation of the Healthy Neonatal Brain into 107 Regions Using Atlas Propagation through Intermediate Time Points in Childhood. Front Neurosci 2016, 10, doi:10.3389/fnins.2016.00220.

11. Moehrlen, U.; Ochsenbein-Kölble, N.; Stricker, S.; Moehrlen, T.; Mazzone, L.; Krähenmann, F.; Vonzun, L.; Zimmermann, R.; Meuli, M. Prenatal Spina Bifida Repair: Defendable Trespassing of MOMS Criteria Results in Commendable Personalized Medicine. Fetal Diagn Ther 2023, 50, 454–463, doi:10.1159/000533181.

12. Moehrlen, U.; Ochsenbein-Kölble, N.; Mazzone, L.; Kraehenmann, F.; Hüsler, M.; Casanova, B.; Biro, P.; Wille, D.; Latal, B.; Scheer, I.; et al. Benchmarking against the MOMS Trial: Zurich Results of Open Fetal Surgery for Spina Bifida. Fetal Diagn Ther 2020, 47, 91–97, doi:10.1159/000500049.

13. Kuklisova-Murgasova, M.; Quaghebeur, G.; Rutherford, M.A.; Hajnal, J. V.; Schnabel, J.A. Reconstruction of Fetal Brain MRI with Intensity Matching and Complete Outlier Removal. Med Image Anal 2012, 16, 1550–1564, doi:10.1016/j.media.2012.07.004.

14. Makropoulos, A.; Robinson, E.C.; Schuh, A.; Wright, R.; Fitzgibbon, S.; Bozek, J.; Counsell, S.J.; Steinweg, J.; Vecchiato, K.; Passerat-Palmbach, J.; et al. The Developing Human Connectome Project: A Minimal Processing Pipeline for Neonatal Cortical Surface Reconstruction. Neuroimage 2018, 173, 88–112, doi:10.1016/j.neuroimage.2018.01.054.

15. Woolrich, M.W.; Jbabdi, S.; Patenaude, B.; Chappell, M.; Makni, S.; Behrens, T.; Beckmann, C.; Jenkinson, M.; Smith, S.M. Bayesian Analysis of Neuroimaging Data in FSL. Neuroimage 2009, 45, S173–S186, doi:10.1016/j.neuroimage.2008.10.055.

16. Gousias, I.S.; Edwards, A.D.; Rutherford, M.A.; Counsell, S.J.; Hajnal, J. V.; Rueckert, D.; Hammers, A. Magnetic Resonance Imaging of the Newborn Brain: Manual Segmentation of Labelled Atlases in Term-Born and Preterm Infants. Neuroimage 2012, 62, 1499–1509, doi:10.1016/j.neuroimage.2012.05.083.

17. Avants, B.; Epstein, C.; Grossman, M.; Gee, J. Symmetric Diffeomorphic Image Registration with Cross-Correlation: Evaluating Automated Labeling of Elderly and Neurodegenerative Brain. Med Image Anal 2008, 12, 26–41, doi:10.1016/j.media.2007.06.004.

18. Avants, B.B.; Tustison, N.J.; Song, G.; Cook, P.A.; Klein, A.; Gee, J.C. A Reproducible Evaluation of ANTs Similarity Metric Performance in Brain Image Registration. Neuroimage 2011, 54, 2033–2044, doi:10.1016/j.neuroimage.2010.09.025.

19. Smith, S.M.; Jenkinson, M.; Johansen-Berg, H.; Rueckert, D.; Nichols, T.E.; Mackay, C.E.; Watkins, K.E.; Ciccarelli, O.; Cader, M.Z.; Matthews, P.M.; et al. Tract-Based Spatial Statistics: Voxelwise Analysis of Multi-Subject Diffusion Data. Neuroimage 2006, 31, 1487–1505, doi:10.1016/j.neuroimage.2006.02.024.

20. Fedorov, A.; Beichel, R.; Kalpathy-Cramer, J.; Finet, J.; Fillion-Robin, J.-C.; Pujol, S.; Bauer, C.; Jennings, D.; Fennessy, F.; Sonka, M.; et al. 3D Slicer as an Image Computing Platform for the Quantitative Imaging Network. Magn Reson Imaging 2012, 30, 1323–1341, doi:10.1016/j.mri.2012.05.001.

